# Flanking sequence preference modulates *de novo* DNA methylation in the mouse genome

**DOI:** 10.1101/2020.06.12.147991

**Authors:** Izaskun Mallona, Ioana Mariuca Ilie, Massimiliano Manzo, Amedeo Caflisch, Tuncay Baubec

## Abstract

Mammalian *de novo* DNA methyltransferases (DNMT) are responsible for the establishment of cell-type-specific DNA methylation in healthy and diseased tissues. Through genome-wide analysis of *de novo* methylation activity in murine stem cells we uncover that DNMT3A prefers to methylate CpGs followed by cytosines or thymines, while DNMT3B predominantly methylates CpGs followed by guanines or adenines. These signatures are further observed at non-CpG sites, resembling methylation context observed in specialised cell types, including neurons and oocytes. We further show that these preferences are not mediated by the differential recruitment of the two *de novo* DNMTs to the genome but are resulting from structural differences in their catalytic domains. Molecular dynamics simulations suggest that, in case of DNMT3A, the preference is due to favourable polar interactions between the flexible Arg836 side chain and the guanine that base-pairs with the cytosine following the CpG. This context-dependent *de novo* DNA methylation provides additional insights into the complex regulation of methylation patterns in different cell types.

## Introduction

DNA methylation plays important roles during mammalian development and the perturbation of this mark is often associated with human disease. In mammals, DNA methylation is deposited by the *de novo* DNA methyltransferases DNMT3A and DNMT3B, while during replication, the maintenance methyltransferase DNMT1 ensures correct propagation of the methyl mark (1, 2). Numerous genome-wide studies identified the exact position and tissue-specific dynamics of individual methyl groups on DNA. These revealed that the majority of CpGs in mammalian genomes are fully methylated, with the exception of active promoters and cell-type-specific enhancer elements (3, 4). In addition to CpG methylation, non-CpG (or CpH) methylation has been identified in numerous tissues (3, 5, 6), with highest levels found in brain where it is suggested to potentially contribute to gene regulation and neuronal function through readout by the methyl-CpG-binding protein MeCP2 (7–10).

Nevertheless, the mechanisms governing the precise deposition of DNA methylation to the genome remain to be fully understood. Although the mammalian genome is almost entirely methylated (3, 4), several regions rely on active recruitment of *de novo* DNA methylation activity, including promoters, enhancers and CpG islands (11–13), repetitive elements (14, 15), and transcribed gene bodies (16, 17). These sites also show tissue-specific variability of DNA methylation and display altered methylation patterns in many diseases (4, 9, 18–21). These observations suggest that pathways that recruit *de novo* methylation vary from cell type to cell type and that the genome-wide methylation patterns are a composite picture resulting from combined activity of multiple DNMT targeting mechanisms and mechanisms that actively or passively remove methylation. In previous work, we and others have shown that the *de novo* DNMTs associate with genomic regions that are marked by distinct chromatin modifications, resulting in enhanced deposition of methyl marks at these sites. For example, readout of H3K36me3 by DNMT3B targets DNA methylation to transcribed gene bodies (17, 22), while DNMT3A shows increased preference for Polycomb-target sites and H3K36me2 (23–25).

Besides these differences in chromatin-dependent targeting of the *de novo* DNMT enzymes, biochemical studies suggest that DNMT3A and DNMT3B differ in their preference for CpGs based on flanking sequences. In *in vitro* methylation assays, DNMT3A has been found to show preferences towards CpGs flanked by pyrimidines (Y) at the +2 position, downstream of the methylated C, while studies using episomal constructs in cells indicate that DNMT3B shows higher methylation activity at sequences containing purines (R) at the same position (26–31). However, given the complex interactions of DNMTs with chromatin and their site-dependent recruitment, it remains unclear if CpG-flanking preferences are observed genome-wide. Some indications come from analysis of CpH methylation in various tissues. Non-CpG methylation is predominantly deposited by the *de novo* DNMTs (5, 32–35) and tissues that have a predominant activity of either DNMT3A or DNMT3B indicate different CpH methylation flanking preferences. In tissues with high DNMT3A activity, like neurons or oocytes, the predominant CpH methylation motif occurs at CpApC, while in ES cells CpH methylation is predominantly deposited by DNMT3B and occurs at CpApG sites (3, 21, 35–37). Taken together, these results suggest that enzymatic activities of the *de novo* DNMTs could be guided by local sequence context to shape cell-type-specific methylomes.

Here we systematically interrogated the sequence preference of the *de novo* DNA methyltransferases DNMT3A and DNMT3B *in vivo*. By re-evaluating available whole genome bisulphite datasets from cells expressing *de novo* DNMTs in *Dnmt*-triple-KO mouse embryonic stem cells lacking DNA methylation, we measured how sequence context influences deposition of methylation at CpG and CpH sites, genome-wide. We identify similarities and enzyme-specific preferences of methylation activity at cytosines in different sequence contexts. We show that DNMT3A prefers to methylate cytosines at TA<u>C</u>GYC sites while DNMT3B prefers to methylate TA<u>C</u>GRC. The same downstream preference is retained at non-CpG sites, with pronounced preferences for TA<u>C</u>ACC and TA<u>C</u>AGC, respectively. Through analyzing the genome-wide distribution of flanking sequence preferences, and furthermore, by investigating methylation in cells with altered targeting of DNMT3A or DNMT3B, we show that these preferences are largely independent of the genomic location or chromatin-mediated recruitment of DNMTs. The analysis of available crystal structures and atomistic simulations of the complex between the DNMT3A catalytic domain and a DNA duplex, suggest potential mechanisms underlying this specificity.

## Results

### *De novo* DNA methyltransferases display flanking sequence preferences in murine stem cells

To examine DNA sequence specificities of *de novo* methyltransferases *in vivo*, we first made use of available whole-genome bisulphite sequencing (WGBS) from *Dnmt*-triple-KO (TKO) mouse embryonic stem cells where the *de novo* DNA methyltransferases DNMT3A2 or DNMT3B1 were reintroduced to the genome (17). In these cells, replication-coupled maintenance of *de novo* methylated DNA is absent without DNMT1, and the WGBS data directly allows us to identify *de novo* DNA methylation activity by DNMT3A or DNMT3B. We leveraged this to investigate if favoured and disfavoured DNA sequence contexts identified reported *in vitro,* are indeed also observed in cells and genome-wide. First, we calculated the average DNA methylation in the TKO-DNMT3A2 and TKO-DNMT3B samples at CpGs in all four CpGpN contexts. Although each CpGpN sequence context was equally represented in the TKO-DNMT3A2 and TKO-DNMT3B WGBS libraries (Supplementary Figure 1a-c), DNA methylation analysis indicated a sequence-preference which differed between DNMT3A and DNMT3B (Figure 1a). The DNMT3A sample showed increased methylation at cytosines in a CpGpT and CpGpC sequence-context, indicating a preference towards pyrimidines at the +2 position from the methylated cytosine (CGY), and DNMT3B at cytosines in a CpGpA and CpGpG context, containing purines at the +2 position (CGR). To further test if the observed difference could be influenced by the genomic context or different library representation of the queried sequences, we re-calculated the methylation scores only on the same CpGpNs that were covered in both, the TKO-DNMT3A2 and the TKO-DNMT3B libraries (Supplementary Figure 1d), and also on CpGpNs that were at least 80% methylated in wild type ES cells (Supplementary Figure 1e). The latter was necessary to exclude that active promoters and enhancers with elevated levels of DNA methylation turnover due to active DNA de-methylation (3, 4, 38) could have a potential influence on the calculated methylation scores. In both cases, the same sequence preference was observed when using these refined CpG sites (Supplementary Figure 1d-e), suggesting that different sequence composition in the WGBS libraries or genomics localisation is not the cause for the observed differences.

**Figure 1.**
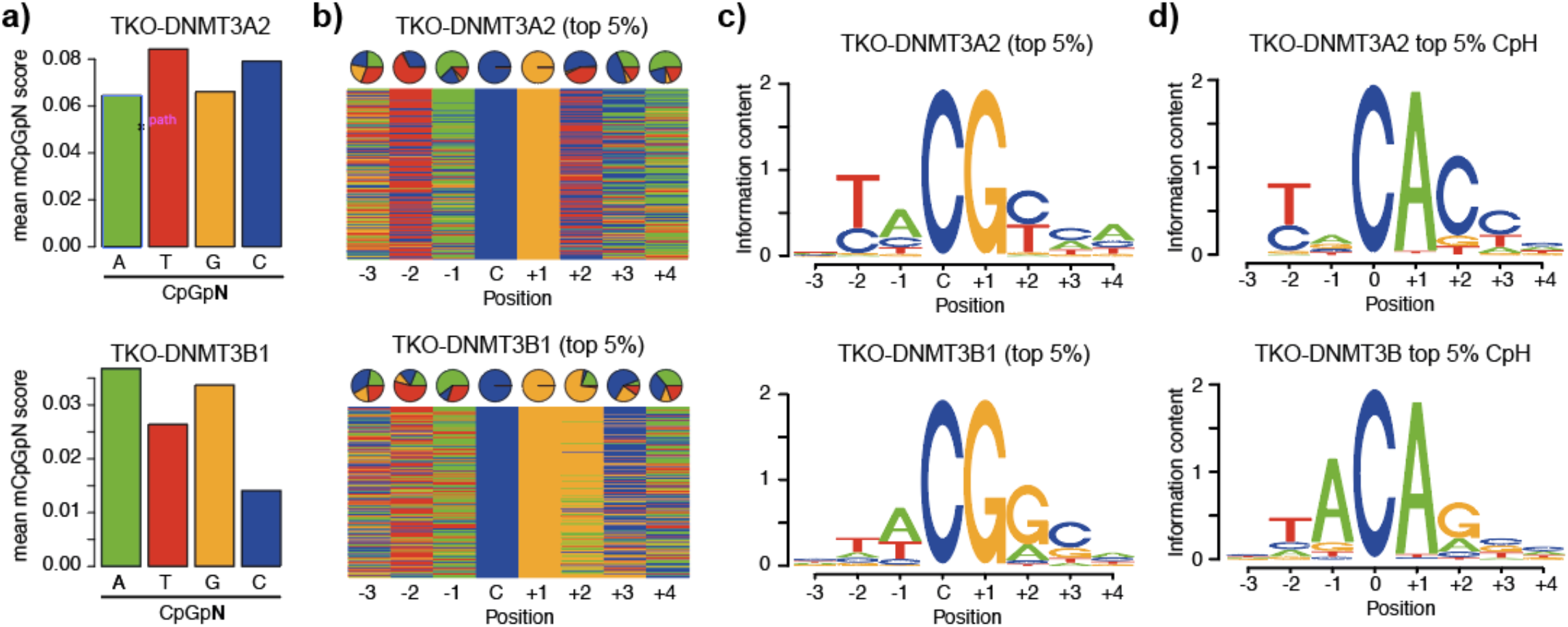
*De novo* DNMTs display flanking sequence preferences at CpG and non-CpG sites. **a)** Bar plots indicating average methylation scores at CpG sites followed by A, T, C and Gs (CpGpN). Average methylation for CpGpNs falling into one of the four categories was calculated from WGBS data obtained from *Dnmt*-TKO ES cells expressing either DNMT3A2 or DNMT3B1. **b)** Nucleotide composition at top 5% from 8-mers ranked by CpG methylation in DNMT3A2 or DNMT3B1-expressing TKO cells. Pie charts show the distribution at the individual positions according to the methylated cytosine: C. **c)** Position weight matrix calculated from the top 5% 8-mers methylated by either DNMT3A2 or DNMT3B indicate the preferred DNA sequence motif. **d)** Position weight matrix calculated from the top 5% 8-mers methylated by either DNMT3A2 or DNMT3B indicate the preferred DNA sequence motif at non-CpG sites.

Following these results, we wanted to have a more detailed view on the sequence preferences of DNMT3A and DNMT3B. Towards this, we expanded our analysis to 8-mer strings with a fixed CpG in the centre (NNNCGNNN) and calculated the methylation scores on the forward or reverse strands separately (Materials and Methods). We first ranked the obtained sequences according to the average methylation score calculated from all 8-mer instances in the *Dnmt*-TKO cells expressing DNMT3A2 or DNMT3B1 (Supplementary Figure 2a-b). This ranking allowed us to identify DNA sequences that were preferentially methylated by DNMT3A and DNMT3B. Top 5%-ranking 8-mers indicated similarities and individual preferences in nucleotides at the queried positions (Figure 1b). For example, we observed that both enzymes have a slight preference for adenines (A) at the −1 position and cytosines (C) at the +3 position. We observed an increased occurrence of thymines (T) at the −2 position at DNMT3B-preferred sites, which was even more prominent at DNMT3A targets (Figure 1b). On top of that, we again observed the +2-nucleotide preference for pyrimidines at DNMT3A targets, where Cs and Ts were equally represented, while DNMT3B displayed an increased preference for purines, especially guanines (G) (Figure 1b). These position-specific preferences flanking CpGs can be further observed when calculating the methylation difference between DNMT3A and DNMT3B for each individual nucleotide position in the 8-mer (Supplementary Figure 2c). Furthermore, these results are in line with previous reports obtained from incubating DNMT3A or DNMT3B with DNA substrates (29–31), indicating that the flanking preferences observed *in vitro* are prevalent in cells and even in presence of chromatin.

By grouping the ranked 8-mers into 20 bins we could furthermore calculate the sequence preferences of each bin and display the results as DNA sequence logos (Figure 1c and Supplementary Figure 3a-b). Importantly we only observed sequence preferences at the top-ranked and bottom-ranked 8-mers, but not at 8-mers ranked in the centre (Supplementary Figure 3a-b). The least methylated 8-mers allowed us to identify sequences that were disfavoured by the enzymes. These predominantly contained purines (G/A) at the −2 position for both enzymes, but we also observed differences at the −1 and +2 position, where DNMT3B showed reduced methylation preference near pyrimidines located at −1 and +2, while DNMT3A did not indicate strong occurrence of disfavoured nucleotides (Supplementary Figure 3a-b). Analysis of sequencing coverage distribution for all 8-mers again results in identical coverages between TKO-DNMT3A2 and TKO-DNMT3B (R=0.995, Supplementary Fig. 4a-b). In addition, we generated an additional WGBS dataset in an independently-derived ES cell clone expressing DNMT3A2 in a newly generated *Dnmt*-TKO background (39). Again, we observed the identical flanking preferences for DNMT3A (Supplementary Figure 4c). Taken together, these results indicate that the observed sequence preferences stem from enzymatic preferences and are not introduced by biases in the WGBS datasets.

Finally, we extended this analysis to all sequences containing a CpH site in the centre of the 8-mer (NNNCHNNN). Ranking all sequences by their average methylation score results in a high preference for CpA methylation for both enzymes (Figure 1d and Supplementary Figure 5a), in accordance with previous results obtained from *in vitro* assays (40) or analysis of non-CpG methylation in mammalian genomes (41). Both enzymes show a preference for T and A at position −2 and −1, respectively (Figure 1d and Supplementary Figure 5a). In addition, we observe again a DNMT3A-specific methylation preference for CpA sequences frequently followed by a C at position +3 (CpApC), while DNMT3B prefers CpApG sites, with position +3 being less defined (Figure 1d and Supplementary Figure 5a-b). Importantly, the observed motifs faithfully resemble the reported non-CpG motifs observed in ES cells or neuronal tissues (3, 6, 21, 41), indicating that these indeed stem from the activity of the individual *de novo* methyltransferases operating in these cell types. We repeated the ranking independently for sites containing CpA, CpT and CpC dinucleotides, resulting in similar DNMT3-specific sequence preferences for position +2 at CpA and CpT sites, while methylation at CpC-containing sequences was similar between both enzymes, likely due to low methylation scores in this CpH context (Supplementary Figure 5c-d).

### Individual and combinatorial influence of flanking nucleotides on DNMT3A and DNMT3B activity

Ranking of k-mers suggested that the sequence context surrounding the CpG/CpH sites has a strong influence on methylation activity. Next, we wondered if coupling of neighbouring nucleotides could have an influence on DNMT3A or DNMT3B activities and we calculated the methylation scores resulting for all possible combinations at the dinucleotides positioned immediately upstream and downstream of the CpGs (Figure 2a). This analysis indicates that for preceding positions (−2 and −1) DNMT3A and DNMT3B mainly prefer to methylate downstream of TpA dinucleotides (Figure 2a). Similar dinucleotide composition preferences can be observed upstream of CpH methylated sites (Supplemental Figure 6a). Analysis of downstream preferences indicates that DNMT3A methylation is preferentially targeted to CpGs followed by CpC or TpC > CpT or TpA > CpA (Figure 2a). In case of DNMT3B, methylation is preferentially targeted to CpG dinucleotides followed by GpC ≫ ApG at positions +2 and +3 (Figure 2a). These downstream di-nucleotide preferences are partially resembled at CpH sites, where DNMT3A prefers to methylate primarily non-CpGs upstream of CpC dinucleotides and DNMT3B sites followed by GpC > GpG or GpT (Supplemental Figure 6a). The results at CpH sites suggest a more defined flanking sequence preference at position +2, where only cytosines and guanines appear to be enzymatic signatures of DNMT3A and DNMT3B, respectively.

**Figure 2.**
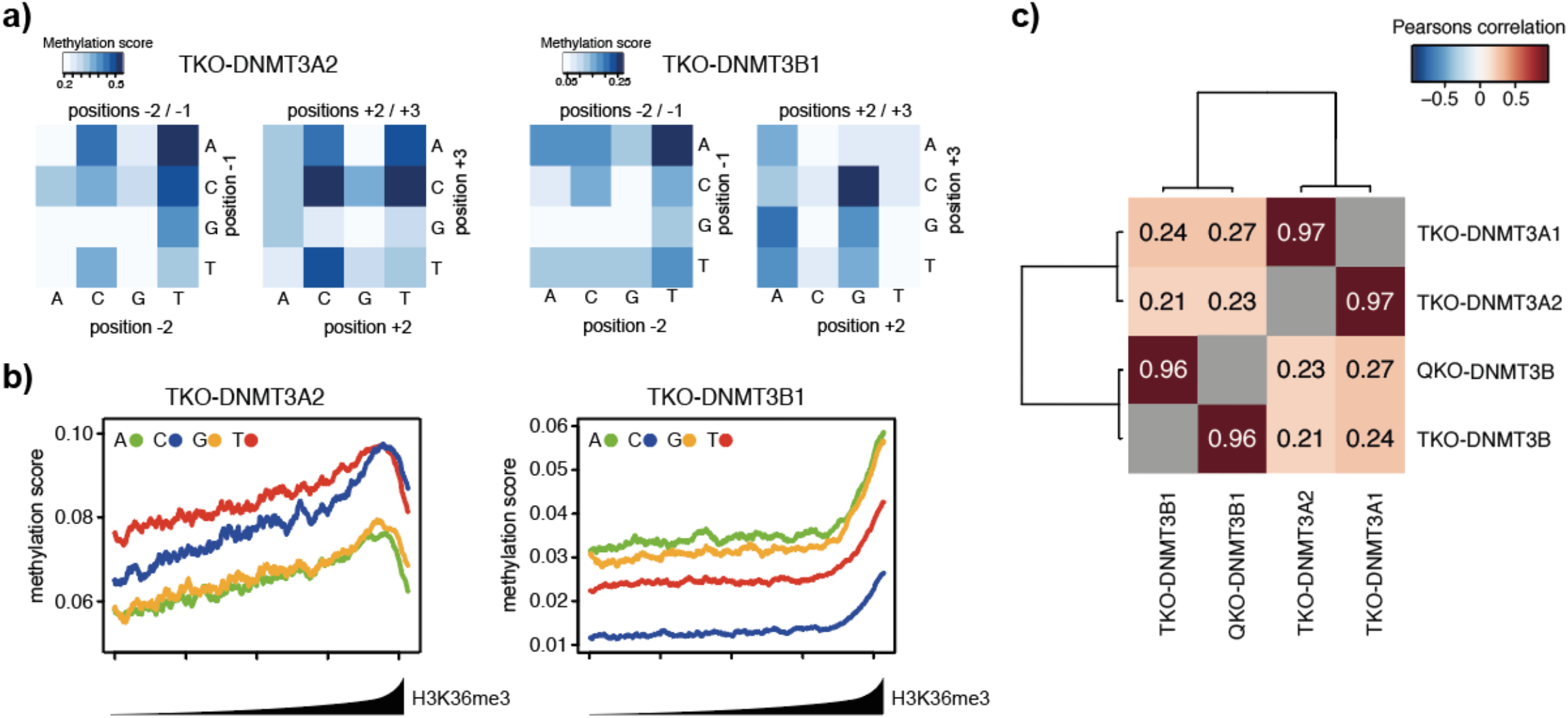
Flanking sequence preferences are independent of the genomic location of the DNMTs. **a)** Heatmap showing the effect of dinucleotide coupling at positions −2/−1 or +2/+3 on CpG methylation (at position 0/+1). **b)** Preferential *de novo* methylation of purines by DNMT3B is not altered by its general preference for H3K36 tri-methylated sites. Shown are de novo DNA methylation at all four CpGpN context genome-wide in relation to H3K36me3 enrichment. 1-kb-sized genomic intervals were ranked and grouped by H3K36me3 enrichment (1000 intervals per bin) and DNA methylation was calculated per bin. Lines indicate mean DNA methylation per bin in TKO cells expressing DNMT3A2 or DNMT3B1. **c)** Heatmap indicating strong correlation in DNA sequence preference between the DNMT3A1 and DNMT3A2 isoforms, or between DNMT3B in presence and absence of H3K36me3. Correlations between identical samples were removed and are shown in grey.

### Differential genomic localisation of DNMTs does not influence sequence preferences

The DNMT3 proteins display distinct genome-wide localisation patterns correlating with the genomic distribution of histone modifications. In case of DNMT3B, methylation is preferentially targeted to H3K36me3 (17). To further investigate if the observed sequence preference is resulting from this differential genomic localisation, we analysed the genome-wide distribution of CpGpN methylation according to H3K36me3 in DNMT3A2 and DNMT3B-expressing *Dnmt*-TKO cells. We ranked 1kb genomic intervals according to H3K36me3 levels and calculated the mean methylation score at CpGpN sites for each interval. While total CpG methylation by DNMT3A2 or DNMT3B varies depending on H3K36me3 levels, as previously reported (17), the enzyme-specific flanking preferences for purines and pyrimidines are not influenced by genomic distribution and their ratios remain constant independent of genomic position (Figure 2b).

To directly test if genomic binding of DNMT3B contributes to the observed flanking preferences, we re-evaluated additional WGBS datasets where we deleted SETD2, the enzyme responsible for deposition of H3K36me3 (42), in *Dnmt*-TKO cells expressing DNMT3B (here termed quadruple-KO, QKO). We and others have previously shown by ChIP-seq, that in absence of SETD2, binding and methylation activity of DNMT3B to sites previously decorated by H3K36me3 is lost (17, 22). This is also observed when we reanalyse the DNA methylation in the context of CpGpN, where all four sequence contexts are equally affected but retain the preference of purines over pyrimidines (Supplementary Figure 7a). By further comparing the methylation scores at 8-mers obtained from DNMT3B-QKO cells with cells containing H3K36me3, we observe the same flanking preference for CpG and non-CpG sites, despite the lack of de novo methylation targeting to H3K36me3 sites (Figure 2c and Supplementary Figure 7b-e). In addition, we have investigated if the previously-reported differential genomic localization of DNMT3A isoforms 1 and 2 (23, 24) would influence flanking sequence preferences. By comparing the results obtained in TKO-DNMT3A2 with a WGBS dataset obtained from *Dnmt*-TKO cells expressing the longer DNMT3A1 isoform, we could not observe any differences in methylated sequence contexts (Figure 2c and Supplementary Figure 7g-i). Taken together, these independent genome-wide comparisons of different targeting sites and flanking-sequence preferences highlight that differential genomic localisation of the *de novo* DNMTs is not the cause for the observed sequence preferences (Figure 2 and Supplementary Figure 7j). While differential genomic localisation of DNMTs indeed influences targeting of DNA methylation, this does not influence the ratio of purines versus pyrimidines at the targeted sites, which remains constant along the genome. These results suggest enzymatic differences between DNMT3A and DNMT3B, rather than genomic localisation, as the source of the observed flanking sequence preference.

### Direct comparison between DNMT3A and DNMT3B reveals structural constraints on methylation preference

The observed sequence preferences could stem from structural differences in the catalytic domains of DNMT3A and DNMT3B. Towards this we first focused our attention on divergent amino acids in the catalytic domains of DNMT3A and DNMT3B. Of the 279 residues in the catalytic domain 44 vary between DNMT3A and DNMT3B (Supplementary Fig. 8a). We made use of available crystal structures of DNMT3A in complex with DNA (43) (PDB ID:6F57) and used DNAproDB (44) with standard interaction criteria to identify which of the variant amino acids interact with the DNA upstream and downstream of the methylated CpG (Figure 3a). We observe that only two variant amino acids are in close contact with the nucleotide bases around the methylated CpG (Figure 3a and Supplementary Fig. 8a). The catalytic loop residue I715 (N656 in DNMT3B) sits in the minor groove of the DNA and contacts the G at −1 position on the same strand, and the A at the −2’ position on the opposite strand of the modified CpG (Figure 3a-b and Supplementary Fig. 8b). The corresponding N656 in DNMT3B crystal structures (45) shows a similar interaction with the minor groove and contacts the A at the −2’ position on the opposite strand (Figure 3a-b and Supplementary Fig. 8c). The interactions of I715 and N656 with the adenine at position −2’ could contribute to the elevated preference for Ts at position −2 on the methylated strand, which is observed for both enzymes (Figures 1c-d).

**Figure 3.**
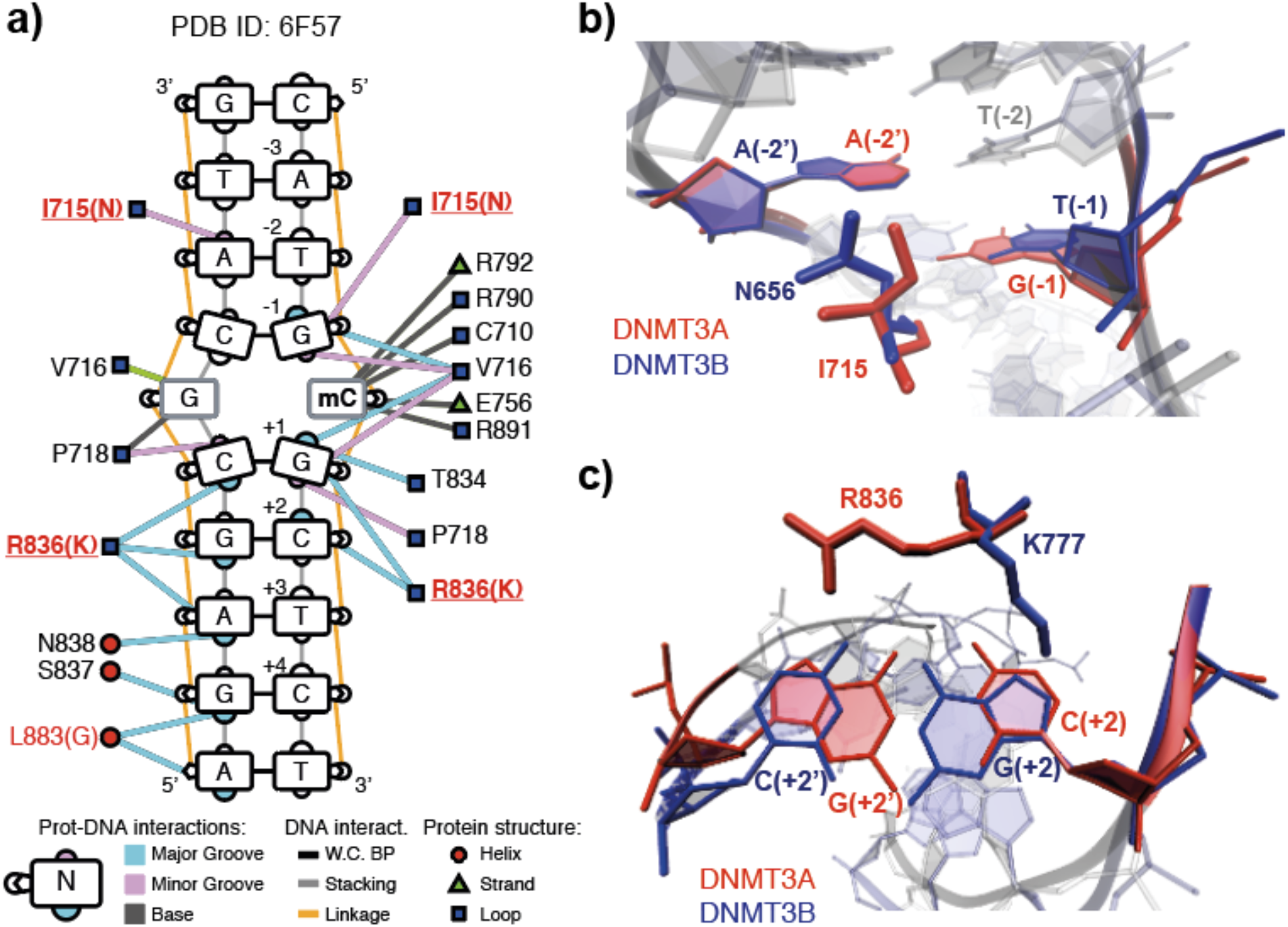
Structural differences between DNMT3A and DNMT3B provide potential cues for observed sequence preference. **a)** Schematic overview of DNAproDB-reported intermolecular interactions between DNMT3A and DNA, based on PDB ID: 6F57. Edges represent interactions between DNMT3A amino acid residues and DNA, while colours represent interactions with the major groove in blue, minor groove in pink and bases in grey. Position of interacting amino acids within DNMT3 protein secondary structures are shown as red circles for alpha helices, green triangles for beta strands and blue squares for loops. **b)** Residues I715 of DNMT3A (PDB ID: 6F57) and N656 of DNMT3B (PDB ID: 6KDA) interact with the minor groove of the DNA by contacting the A at the −2 position on the opposite strand of the modified CpG. **c)** Residues R836 of DNMT3A and K777 of DNMT3B interact with the DNA downstream of the methylated CpG through interactions with the major groove at position +2. The side chains show opposing orientation, with K777 (PDB ID 6KDA) contacting the guanine on the methylated strand and R836 (PDB ID 6F57) contacting the guanine on the opposite strand.

The variant residue R836 in the target recognition domain (TRD) of DNMT3A (K777 in DNMT3B) contacts the +1 to +3 nucleotide pairs in the major groove downstream of the methylated CpG (Figure 3a). This residue was initially shown to be required for the DNMT3A-specific preference for G at the +1 position in the CpG site (43). Interestingly, in DNMT3B, it is rather N779 (N838 in DNMT3A) that mediates the specificity for the G nucleotide at position +1 of the methylated CpG, while the corresponding K777 contributes towards the DNMT3B-specific flanking preference for Gs at position +2 (31, 45). Given these observations, we wanted to investigate how the DNMT3A R836 contributes to the observed flanking preference at position +2. Superimposition of available structures of DNMT3A and DNMT3B in complex with DNA (43, 45) indicate that both residues (DNMT3A R836 and DNMT3B K777) can contact the nucleotides at position +2, but the side chains are oriented in opposite directions (Figure 3c). In the respective crystal structures, the DNMT3B K777 side chain contacts the guanine at position +2 on the methylated strand, while the DNMT3A R836 side chain contacts the guanine at position +2’ on the opposite strand (Figure 3c and Supplementary Fig. 8d-e). The opposing orientations and interactions on opposite strands could point to a potential mechanism underlying the observed DNMT-specific preference at position +2.

### Atomistic simulations reveal interaction dynamics between key residues and DNA

To investigate the structural plasticity of the main interactions we performed molecular dynamics (MD) simulations starting from a DNMT3A/decameric DNA complex extracted from the crystal structure of DNMT3A-DNMT3L-DNA (PDB ID: 5YX2) (43). Two simulation systems were prepared differing only at the position +2 of the 10-mer DNA duplex, namely CATAmCG**C**CCT and CATAmCG**G**CCT (boldface for position +2, Supplementary Fig. 9a). The nucleotides preceding the methylation site were mutated to T and A to accommodate the observed preferences of DNMT3A and DNMT3B. Both systems contained also a molecule of the co-product S-adenosyl-L-homocysteine (SAH) (Supplementary Figure 9b). Five independent simulations were carried out for each system for a total sampling of 5 μs (Materials and Methods). We focused the analysis on the interaction between R836 and the DNA duplex and in particular the closest nucleotide, that is G or C at position +2’ on the unmethylated strand (Figure 3c) and calculated the shortest distance between any pairs of non-hydrogen atoms along the MD trajectories. In presence of the disfavoured sequence, the histogram of the distance between R836 and the cytosine at position +2’ shows a maximum at about 4 Å which corresponds to van der Waals contacts (Figure 4a and Supplementary Movie 1). Interestingly however, in the presence of the favoured sequence the distance between R836 and the guanine at position +2’ shows a bimodal distribution with a peak at hydrogen bond distance (~ 2.7 Å) and a broad region up to 5.5 Å (Figure 4a). This indicates a potential switch in the orientation of the R836 side chain, resulting in two conformational states which we here call “R836-in” and “R836-out”. Indeed, the simulations reveal that in presence of the preferred sequence the guanidinium group of R836 can form two hydrogen bonds, respectively, with the carbonyl oxygen and N7 nitrogen of the guanine at position +2’ (i.e., the guanine that base-pairs with C at +2) resulting in the “R836-in” state (Figure 4b-c, Movies 2-3, and Supplementary Figures 9d-e). This switching of R836 is in line with a recent crystal structure of DNMT3A in complex with DNA containing the disfavoured CpGpA sequence, suggesting that this side chain dynamically interacts with DNA, depending on the flanking nucleotides downstream of CpG (46). In our simulations, the total interaction energy between the side chain of R836 and its surrounding (DNA, protein, and solvent) is similar for the disfavoured DNA sequence and the two states of the favoured DNA sequence (Supplementary Figure 9c). Concerning the individual contributions, the interaction energy between R836 and DNA is about 15 kcal/mol more favourable for the “R836-in” state than the “R836-out”, and the latter is similar as in the disfavoured sequence (Supplementary Figure 9c). The difference in interaction energy originates almost only from the aforementioned hydrogen bonds. The additional hydrogen bonds with DNA (more precisely with the G at position +2’ in the favoured DNA sequence) are compensated by a loss of interactions between R836 and the solvent (Supplementary Figure 9c). Thus, the MD simulations indicate that the direct interactions between R836 and methylated DNA duplexes containing mCGC sites are more favourable compared to mCGG sites, while the interaction energy of R836 and the rest of the system (protein/DNA complex and solvent) is similar.

**Figure 4.**
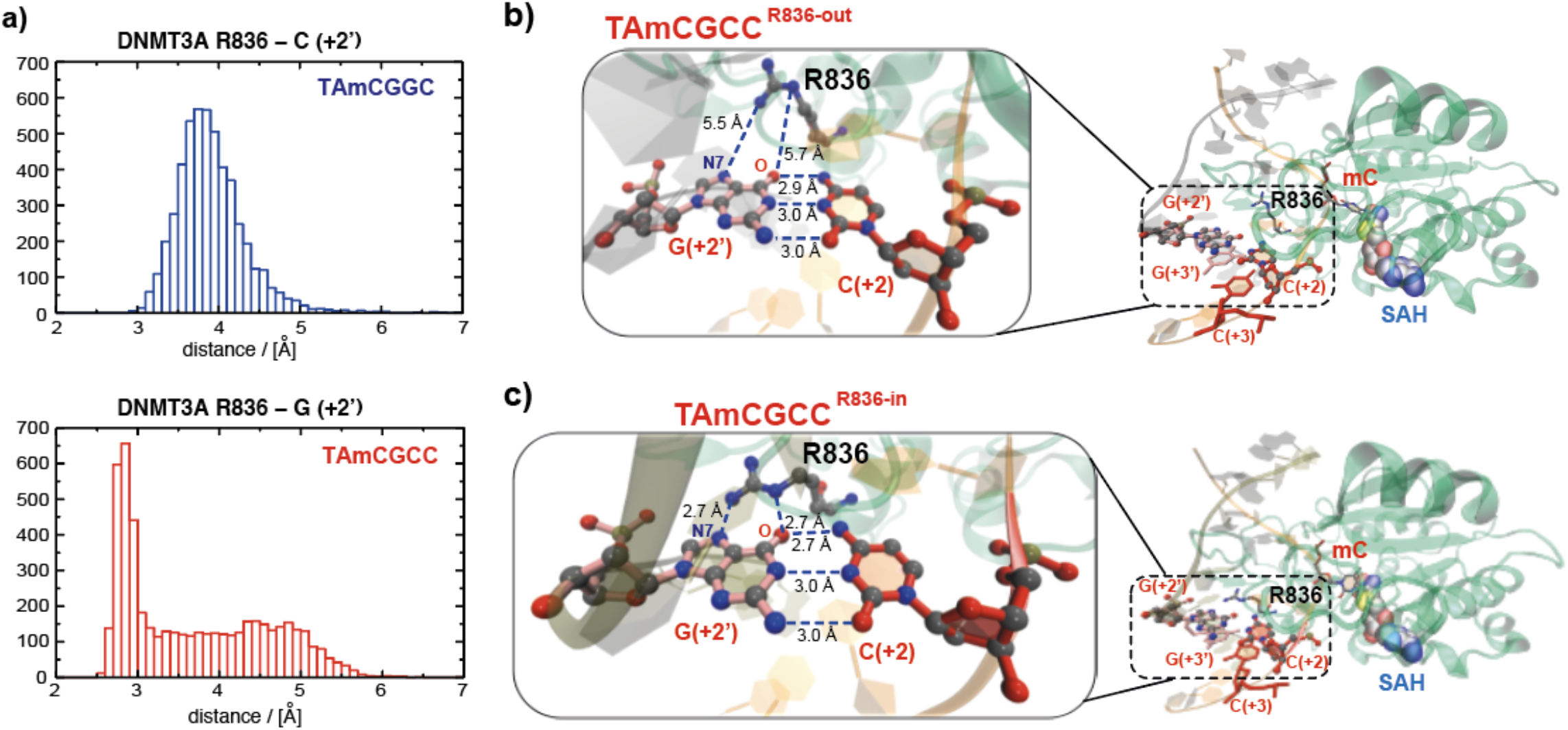
Molecular dynamics simulations reveal potential mechanisms underlying DNMT3A preference for CpGpC sites. **a)** Histograms displaying the measured distance between the R836 side chain and the base at position +2’ for the disfavored (top) and the favored (bottom) DNA duplex sequence. For the favored sequence, a broad region and a peak are observed corresponding to the state before the conformational switch (“R836-out” state) and after (“R836-in” state). **b-c)** The two interactions states of DNMT3A R836 and the preferred DNA sequence obtained from the MD simulations. Magnifications show the distances between DNMT3A R836 and the guanine at position +2’ for the “R836-out” (b) and “R836-in” state (c). The hydrogen bonds between the nitrogen atoms of the guanidinium group of R836 and the carbonyl oxygen and N7 nitrogen of the guanine at position +2’ are indicated.

## Discussion

Sequence-specific flanking preferences have been described for the *de novo* DNA methyltransferases based on in vitro methylation experiments or using episomal constructs in cells (26–28, 30, 31, 47). In addition, analysis of tissue-specific WGBS data indicates that sequence-context of non-CpG methylation strongly varies between tissues expressing DNMT3A or DNMT3B (3, 21, 35–37). Together, these reports suggest that DNA sequence preferences of the *de novo* DNMTs could provide an additional layer of methylome regulation. However, it remained to be clarified if these enzyme-specific preferences occur *in vivo* and genome-wide. Especially, given the reported differential localisation and cell-type-specific activities of *de novo* DNMTs, it remained a challenge to directly and comprehensively compare the enzymatic preferences of these enzymes in the same genomic environment. By re-evaluating whole-genome bisulphite sequencing data obtained from *Dnmt*-TKO ES cells lacking DNA methylation where we have previously re-introduced the individual *de novo* enzymes, we were able to characterize the genomic flanking preferences of DNMT3A and DNMT3B at unprecedented detail. Here we show that DNMT3A prefers to methylate predominantly in a NTA<u>C</u>GYCN context, while DNMT3B prefers NTA<u>C</u>GRCN. We find nucleotides around the methylated site that are equally preferred by both enzymes and furthermore, highlight that the +2 nucleotide following the methylated cytosine is the strongest determinant of enzyme-specific sequence preference, confirming previous *in vitro* measurements. This sequence preference outside of the methylated CpG dinucleotide was largely consistent with methylation preferences observed around non-CpG sites, with NTA<u>C</u>ACCN and NTA<u>C</u>AGCN being the most-favoured sites for DNMT3A and DNMT3B, respectively. The non-CpG-specific flanking preferences are identical to non-CpG motifs reported in tissues where either DNMT3A predominantly contributes to *de novo* methylation, such as neurons or oocytes, or in ES cells, where DNMT3B seems to contribute to the majority of non-CpG methylation (3, 6, 21, 41) – confirming that the methylation patterns observed in these cells are indeed a footprint of enzyme-specific *de novo* DNMT activities.

We furthermore show here that the observed sequence preference does not stem from the differential localisation of the *de novo* DNMTs to the genome but can be attributed to minor variations in the structure of the TRD loop in the catalytic domain. Direct comparison of protein sequences and structures of DNMT3A (43) or DNMT3B (45) together with DNA indicates a variant residue that contacts position +2 downstream of the methylated CpG in the major groove of the DNA. The arginine side chain of this residue in DNMT3A (R836) contacts purines on the opposite strand from the methylated CpG, while a lysine in DNMT3B at the same position (K777) contacts purines on the methylated strand. The latter interaction was recently shown to influence the flanking preference of DNMT3B for CpGpG sites *in vitro* (31, 45), suggesting that R836 could play similar roles in determining DNMT3A preferences for purines on the opposing strand, resulting in the observed CpGpY flanking preference on the methylated strand. To shed light on the atomistic details of the DNMT3A/DNA duplex interactions we performed MD simulations with decameric DNA products containing favoured and disfavoured flanking sequences (difference at position +2). The quantification of interaction distances along the MD trajectories provides information that goes beyond a simple interpretation of the crystal structures. The favoured DNA sequence and R836 display two conformational states “R836-in” and “R836-out”. In the first state stabilizing hydrogen bonds between the side chain of R836 and the G that base-pairs with the C at position +2 of the favoured sequence are formed, whereas in the latter only weaker van der Waals contacts are observed. This dynamic switching of R836 is fully consistent with recent crystal structures that describe flexible conformations of the R836 side chain, depending on the flanking sequence (43, 46). Together with our molecular dynamics simulations, these results support a sequence-dependent conformation of the R836 side chain, and pinpoint R836 a critical residue in mediating the flanking sequence preferences observed for DNMT3A.

While our simulation results seem intuitive at first, one has to mention that stronger interactions with the preferred sequence could hinder DNA dissociation after the methylation reaction, and the “R836-out” state with weaker interactions could potentially be more important for the observed sequence preference. Indeed, product release has been shown to be the rate-limiting step regulating the turnover rate of the paradigm bacterial DNA methyltransferase M.*HhaI* (48), and DNA adenine-methyltransferases (49, 50) and future experiments are required to investigate if the same principle applies to mammalian *de novo* DNMTs. In general, our MD simulations have revealed the atomistic details and intrinsic flexibility of the DNMT3A/DNA duplex interactions. Because of the accessible time scales, which are in the μs range it is not possible to speculate on the influence of individual interactions on the association kinetics of the enzyme/substrate complex and dissociation kinetics of the product.

Understanding the sequence-context-dependent methylation activities of DNMT3A and DNMT3B in living cells provides novel insights into how DNA sequence could shape DNA methylation patterns. We suggest that sequence-dependent *de novo* methylation provides an additional regulatory layer of the methylome landscape with potential impact on gene activity through recognition of the methylated cytosine in different sequence contexts. This is for instance exemplified by MeCP2, a methyl-binding protein with *in vitro* and *in vivo* binding preference for methylated CpApC and to some extent CpApT sites (7). Methylated CpApC sites drive recruitment of this protein in neuronal cells and lack of DNMT3A has been shown to disrupt this localization, resulting in changes of gene expression relevant for neuronal function (7, 51). In addition, several transcription factors (TFs) have been shown to be influenced by the presence of DNA methylation in their recognition motifs (39, 52, 53), and DNMT-specific methylation signatures are expected to differentially influence genomic binding of TFs to their cognate sites. Loss and gain of function mutations in the *de novo* DNA methyltransferases and the corresponding changes in DNA methylation patterns have been associated with numerous diseases and cancers, with DNMT3A mutations being highly prevalent in hematological malignancies (54, 55). Interestingly, R882H, one of the frequent DNMT3A mutations in acute myeloid leukemia patients has been found to switch the flanking sequence preference of DNMT3A to resemble DNMT3B *in vitro* (29, 30, 46, 56). Future studies will help to integrate cell-type-specific activities of DNMTs, deposition and readout of sequence-specific methylation patterns in order to understand how these contribute to gene activity and cellular function in healthy and diseased tissues.

## Acknowledgements

We thank Mark D. Robinson (UZH, DMLS), the Swiss National Supercomputing Centre and S3IT at the University of Zurich for providing the computational infrastructure. We thank Ino Karemaker and Stefan Butz for their critical input on the manuscript. The authors would like to acknowledge following support: Swiss National Science Foundation – SNSF Professorships #157488 and SNSF Spark #190378 to T.B.; SNSF Sinergia #180345 to T.B. and A.C.; SNSF Excellence grant #310030B-189363 to A.C.; I.M.I. thanks the Peter und Traudl Engelhorn Foundation for a postdoctoral fellowship.

## Materials and Methods

### Cell line generation and cultivation

*Dnmt1,3a,3b*-triple-KO cells were obtained from (39). Cells were cultured on 0.2% gelatine-coated dishes in DMEM (Invitrogen) supplemented with 15% fetal calf serum (Invitrogen), 1× non-essential amino acids (Invitrogen), 1mM L-glutamine, leukemia inhibitory factor, and 0.001% b-mercaptoethanol. DNMT3A2 expressing *Dnmt*-TKO cell lines were obtained by recombinase-mediated cassette exchange (RMCE) using pL1-CAGGS-bio-DNMT3A2-polyA-1L as previously described (17).

### Whole-genome bisulphite library preparation and sequencing

Whole-genome bisulphite sequencing for TKO cells expressing DNMT3A2 was performed as described (17). In brief, 6 ug sonicated, genomic DNA were pre-mixed with *SssI*-methylated phage T7 and unmethylated phage Lambda DNA (both 10 ng, sonicated) and sequencing libraries were prepared using the NEB ULTRA kit following manufacturer’s instructions for genomic DNA library construction and using methylated adaptors (NEB <u>E7535S</u>). Adaptor-ligated DNA was isolated by 2% agarose gel electrophoresis (350–400 bp), then converted by sodium bisulphite using the Qiagen Epitect bisulphite conversion kit. Converted libraries were enriched by 10 cycles of PCR with the following reaction composition: 1 μl Pfu TurboCx Hotstart DNA polymerase (Stratagene), 5 μl PfuTurbo Cx reaction buffer, 25 μM dNTPs, 1 μl universal and 1 μl index primer. PCR cycling parameters were: 95C for 2 min, 98C for 30 sec, then 10 cycles of 98C for 15 sec, 65C for 30 sec and 72C for 3 min, ending with one 72 C for 5 min step. The reaction products were purified twice using Ampure-XT beads. Quality of the libraries and size distribution were assessed on an Agilent High Sensitivity tape station. Libraries were sequenced on Illumina HiSeq 4000 machines. Data has been deposited to Gene Expression Omnibus (GEO), under accession number: GSE151992.

### Whole-genome bisulphite data processing

Published WGBS datasets for TKO-DNMT3A2, TKO-DNMT3B1 and QKO-DNMT3B1 obtained from GSE57411, and newly-generated WGBS data (TKO-DNMT3A2_r2) were pre-processed before alignment to the mouse genome. For bulk analysis of CpGpN sites, the sequencing samples were processed and aligned as described in(17). In brief, reads were first trimmed to 50nt using fastx_trimmer and then aligned to the mouse genome (mm9) using BOWTIE in QuasR qAlign() with standard options and bisulfite=“undir” setting(57). Methylation calls for CpGs were obtained using the qMeth() function in QuasR in “CpGcomb” mode and excluding CpGs overlapping with known SNPs. Bisulphite conversion efficiency was measured by spiked-in controls of methylated T7 DNA and unmethylated lambda DNA.

For detailed WGBS analysis using 8-mer motifs, sequencing reads were pre-processed and trimmed using cutadapt v1.16 (58) and sickle v1.33 (Joshi and Fass 2011, https://github.com/najoshi/sickle). Read quality was checked before and after processing using FastQC v0.11.5 (https://www.bioinformatics.babraham.ac.uk/projects/fastqc/). Trimmed reads were mapped against the mm9 mouse assembly using bwa-meth v0.2.0 (59) running bwa v0.7.12-r1039 (60). Mapping quality was evaluated with qualimap v2.2.1 (61). Duplicates were removed with Picard MarkDuplicates (picard-tools v1.96, http://broadinstitute.github.io/picard/) with ‘REMOVE_DUPLICATES=TRUE and REMOVE_SEQUENCING_DUPLICATES=TRUE’ flags. Alignments with MAPQ > 40 were used as input to methylation calling with methyldackel v0.3.0-3-g084d926 (https://github.com/dpryan79/MethylDackel) (using HTSlib v1.2.1) in ‘cytosine_report’ mode for all cytosines in the genome both CpG and CpH contexts (‘extract -q 40 --cytosine_report --CHH – CHG’).

### Sequence-dependent DNA methylation analysis

For global analysis of methylation preferences at CpGpN sites, we first calculated the methylation scores for individual CpG instances with min coverage = 10 and max coverage = 50 in the WGBS datasets. Methylation scores were obtained as: number of methylated reads / (number of methylated +unmethylated reads) per CpG. We obtained the CpGpN sequence context by extending the CpGs by one nucleotide downstream on the + strand and querying the mm9 sequence using the biostrings package in R. Average methylation scores were calculated for all CpGs falling in one of the four contexts: CpGpA, CpGpT, CpGpG or CpGpC. The same analysis was performed using only CpGpNs that were only covered in both datasets according to the criteria described above (TKO-DNMT3A2 and TKO-DNMT3B1) and CpGpN sites that are at least 80% methylated in wild type ES cells. To evaluate the genome-wide distribution of CpGpN methylation according to H3K36me3, the mouse genome was partitioned into 1kb, nonoverlapping intervals using GenomicRanges in R. Intervals overlapping with satellite repeats (Repeatmasker), ENCODE black-listed and low mappability scores (below 0.5) were removed in order to reduce artefacts due to annotation errors and repetitiveness. First, we calculated the signal strength of H3K36me3 at each individual interval by counting the numbers of reads from available ChIP-seq experiments (62). All genomic intervals were first ranked based on the total number of H3K36me3 ChIP-seq reads. Then, the ranked intervals were binned into groups of 1000 intervals, resulting in a total of 730 bins. Within each bin, we calculated the average methylation falling into the four possible CpGpN sequence contexts for each WGBS sample.

For the 8-mer specific analysis, first a dictionary containing a complete set of 8-mer motifs for NNNCGNNN and NNNCHNNN was created. Next, we separately counted all methylated or unmethylated WGBS reads aligning in the corresponding orientation to every genomic instance containing the 8-mer of interest and calculated the methylation score. The low-expression of the re-introduced DNMT3 enzymes to TKO cells and the absence of DNMT1-mediated maintenance resulted in sparse methylation, with majority of CpG sites lacking DNA methylation (Supplemental Fig. 2b). To account for the sparse methylation, we defined the methylation status of each strand-specific 8-mer as methylated when at least one methylated read was present. These genomic instances of methylated and unmethylated 8-mers were then collapsed into a list containing total methylated and unmethylated instances per motif. This was done for each WGBS dataset individually, resulting in a data frame summarising methylated (M) and unmethylated (U) reads per motif. Methylation scores per motif were obtained by calculating M/(M+U) and were used for generating motif ranks. Position weight matrices were calculated and displayed using the seqLogo package in R. Nucleotide coupling analysis upstream and downstream of the methylated target site was calculated by first counting all methylated and unmetylated instances containing NNCX or CXNN 4-mers, where X= is either G or H depending on CpG of CpH context. Finally, the methylation score at all possible combinations at −2/−1 or +2/+3 sites was calculated as described above and represented as heatmaps.

### Molecular dynamics system preparation

The crystal structure of DNMT3A-DNMT3L in complex with double-stranded DNA containing two CpG sites (PDB ID: 5YX2) (43) was used as the structural basis of this study. We reduced the complex to one DNMT3A methyltransferase interacting with a 10-mer DNA, containing one CpG site, and an S-adenosyl-L-homocysteine (SAH) cofactor (Supplementary Fig. 8b). For the simulations, the zebularine in the crystal structure was replaced by a cytosine methylated at C5. Furthermore, four nucleotides in the DNA sequence present in the crystal structure (two nucleotides preceding and two following the methylation site) were mutated to accommodate the observed preferences of DNMT3A and DNMT3B (Supplementary Fig. 8a). The mutation of the nucleotides and the reconstruction of the missing residues in the crystal structure were performed using the Maestro software (Schrödinger, release 2020-1). The methyltransferase DNMT3A, the rest of the DNA chain, the cofactor and the crystal water molecules were kept as in the crystal structure.

### Molecular dynamics simulations

Overall, 34 000 node hours using 72 virtual cores per node, were used to carry out the simulations on Piz Daint at the Swiss National supercomputing Centre. All simulations were carried out using the GROMACS 2018.6 simulation package (63) and the CHARMM36m force field (64). Five independent 1-μs simulations with different initial random velocities were carried out for each system. To reproduce neutral pH conditions, the N-terminus was positively charged, while the C-terminus was negatively charged. Furthermore, the 5’- and 3’-termini in the DNA sequence were capped with phosphate groups and OH groups, respectively. Each system was solvated in a cubic box (edge length of 13.9 nm) with TIP3P water molecules (65) to which 150 mM NaCl were added, including neutralizing counterions. Periodic boundary conditions were applied. Following steepest descents minimization, the systems were equilibrated under constant pressure for 2 ns, with position restraints applied on the heavy atoms of the complexes. Temperature and pressure were maintained constant at 300K and 1 atm, respectively, by using the modified Berendsen thermostat (0.1 ps coupling) (66) and barostat (2 ps coupling) (67). For the production runs, performed in the NVT ensemble, harmonic restraints (force constant K = 1000 kJ/(mol∙nm^2^)) were used for the heavy atoms of the end nucleotides and snapshots were saved every 50 ps. The short-range interactions were cut-off beyond a distance of 1.2 nm and the potential smoothly decays to zero using the Verlet cutoff scheme. The Particle Mesh Ewald (PME) technique (68) with a cubic interpolation order, a real space cut-off of 1.2 nm and a grid spacing of 0.16 nm was employed to compute the long-range interactions. Bond lengths were constrained using a fourth order LINCS algorithm with 2 iterations (69). In all simulations the time step was fixed to 2 fs.

## Supplementary Information

### Supplementary Information Content

**Supplementary Movie 1** – Representative sequence from the MD simulations showing R836 in presence of the disfavoured DNA sequence.

**Supplementary Movie 2** – Representative sequence from the MD simulations showing R836 in presence of the preferred DNA sequence. The “out” state is shown, where R836 does not interact with the G at position 2’.

**Supplementary Movie 3** – Representative sequence from the MD simulations showing R836 in presence of the preferred DNA sequence. The “in” state is shown, where R836 interacts with the G at position 2’ through hydrogen bonds between the nitrogen atoms of the guanidinium group of R836 and the carbonyl oxygen and N7 nitrogen of the guanine.

## Supplementary Figures

**Supplementary Figure 1.**
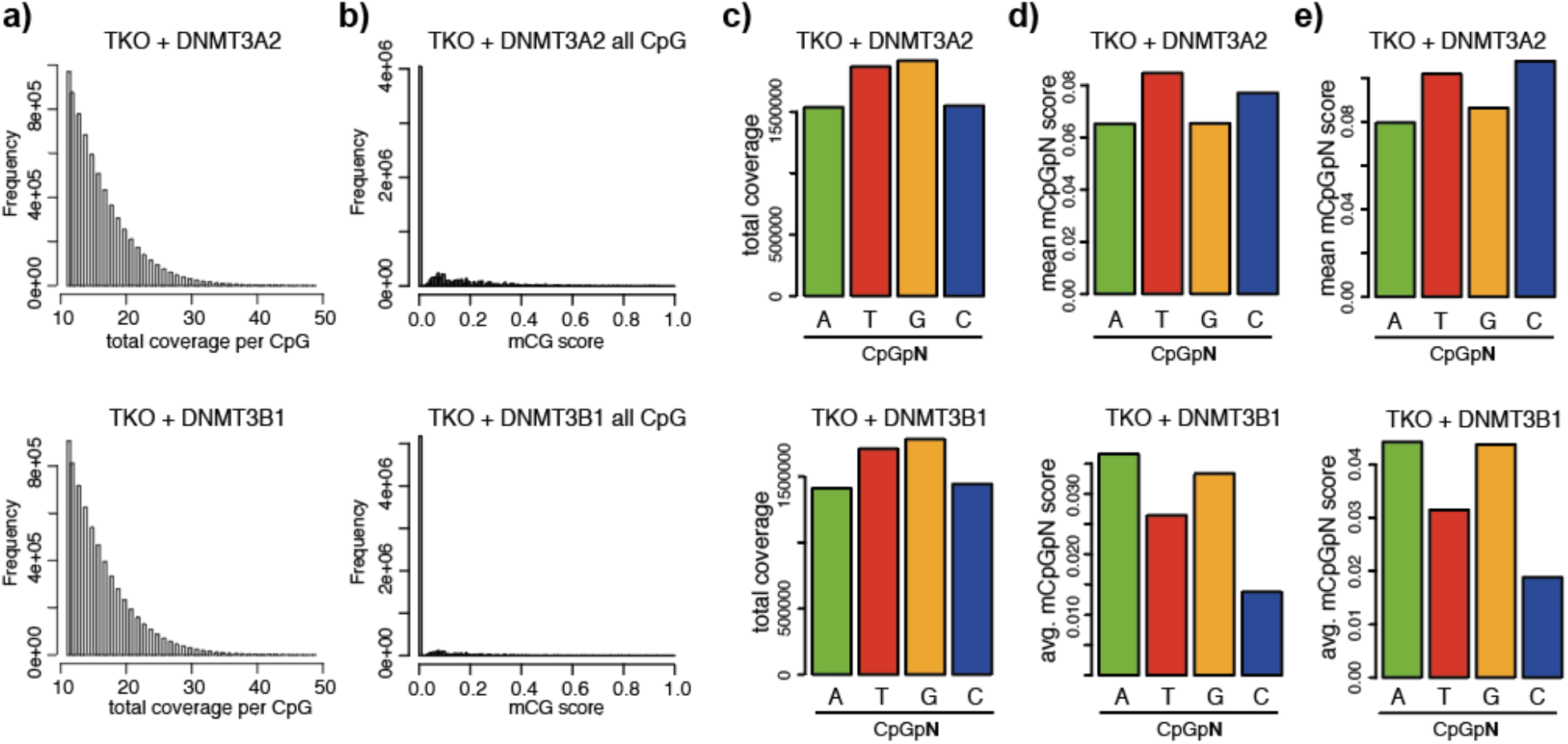
*De novo* DNMTs sequence preferences at CpG sites. **a)** Histogram indicating similar coverage of the analysed CpGs in *Dnmt*-TKO ES cells expressing either DNMT3A2 or DNMT3B1. **b)** Histogram showing the sparse methylation introduced by the individual de novo DNMTs in the *Dnmt*-TKO ES cells. **c)** Bar plots showing the coverage of each CpGpN site analysed in Figure 1a. **d)** Same as Figure 1a but utilising only CpGpN sites that were covered at least 10x in both samples. **e)** Same as Figure 1a but utilising only CpGpN sites that were methylated more than 80% in wild type ES cells.

**Supplementary Figure 2.**
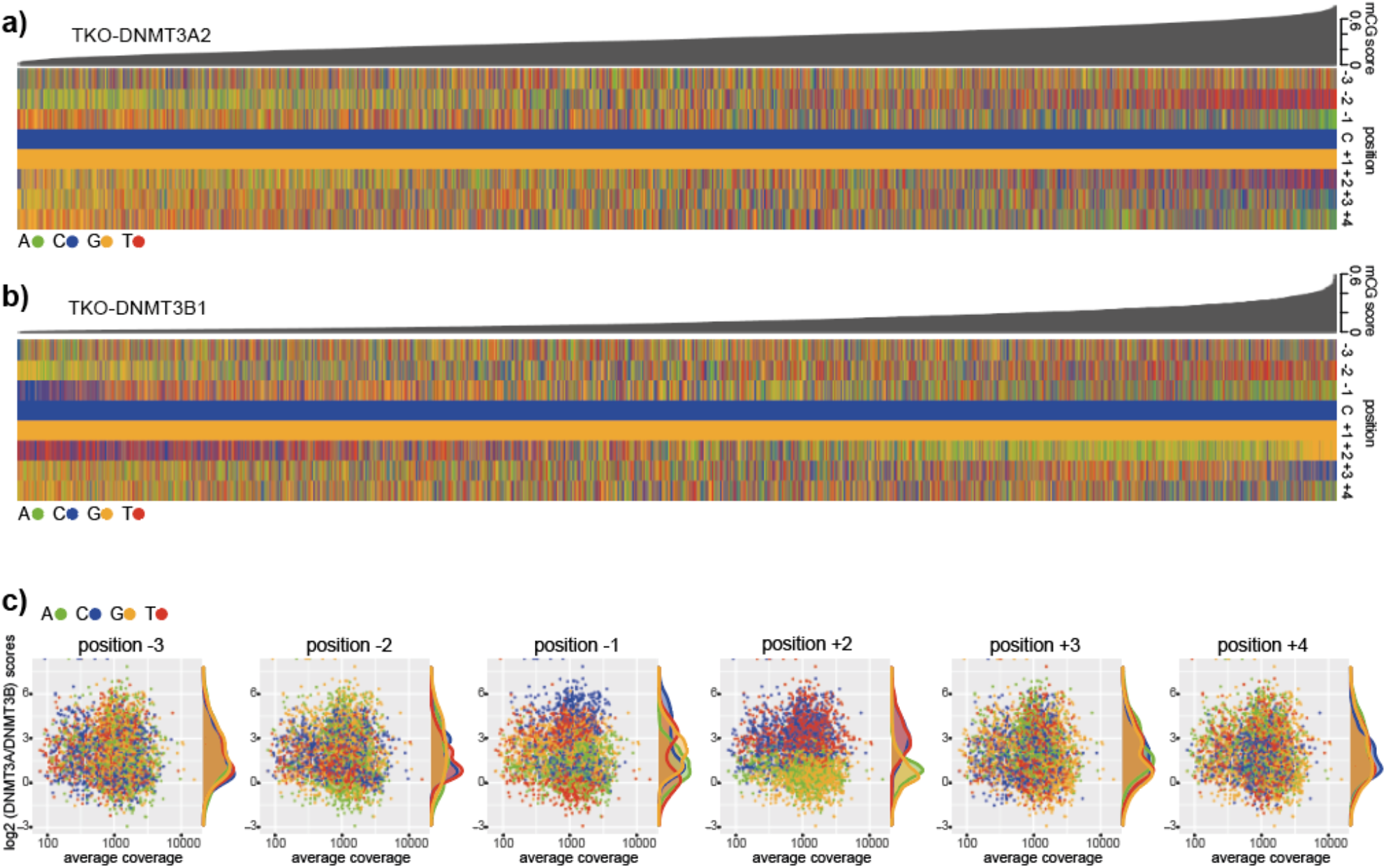
Flanking sequence preferences at CpG sites ranked by methylation score. **a-b)** Heatmap indicating the sequence composition at 8-mers ranked by CpG methylation in TKO-DNMT3A2 or TKO-DNMT3B1. Shown are the nucleotides at positions −2 to +4 around the methylated cytosine. Green: A, blue: C, orange: G and red: T. The average methylation calculated for all instances of each 8-mer is shown above. **c)** MA plots showing log2 methylation differences between DNMT3A2 and DNMT3B1 activity at all analysed CpG 8-mers (y-axis) and the log10 average coverage of each 8-mer (x-axis). All plots show the same data, but the 8-mers are coloured based on the respective nucleotide at positions −3 to +4, excluding static C and G positions 0 and +1.

**Supplementary Figure 3.**
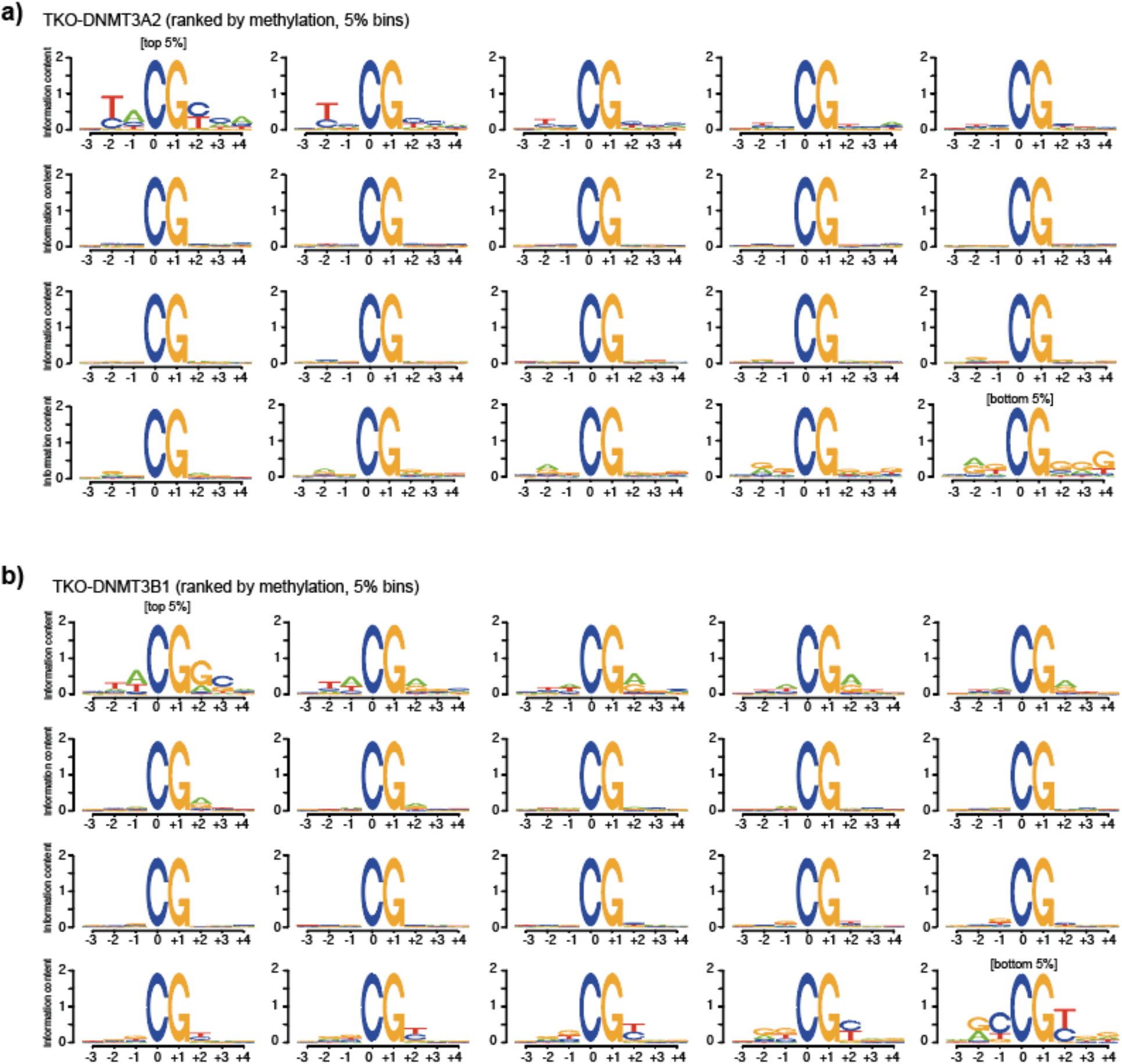
Sequence logos reveal preferred and disfavoured flanking sequences. **a-b)** Sequences ranked by average DNA methylation were collected into 20 bins. Each bin was used to compute the indicated position weight matrix. Shown are bins in decreasing preference from left to right, top down.

**Supplementary Figure 4.**
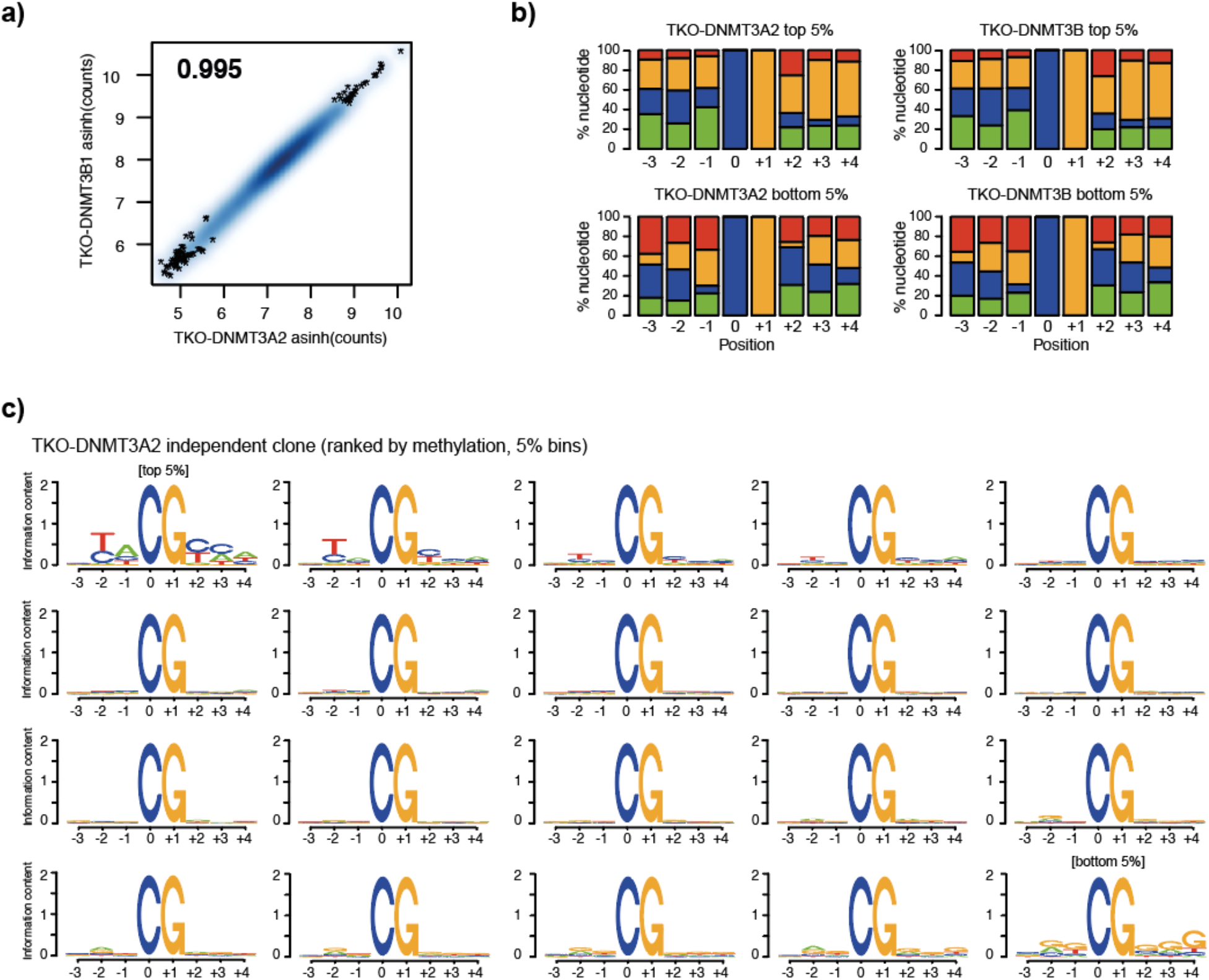
Coverage differences do not influence the observed sequence preferences. **a)** Scatter plot showing similarity in sequence coverage between TKO-DNMT3A2 and TKO-DNMT3B1 samples. Datapoints show the asinh-transformed total reads covering 8-mers containing a CpG in the central position from the WGBS reads. **b)** Stacked bar plots show similarity in sequence composition at 8-mers ranked by read-coverage. **c)** Sequences ranked by average DNA methylation in the TKO-DNMT3A2 replicate were collected into 20 bins. Each bin was used to compute the indicated position weight matrix. Shown are bins in decreasing preference from left to right, top down.

**Supplementary Figure 5.**
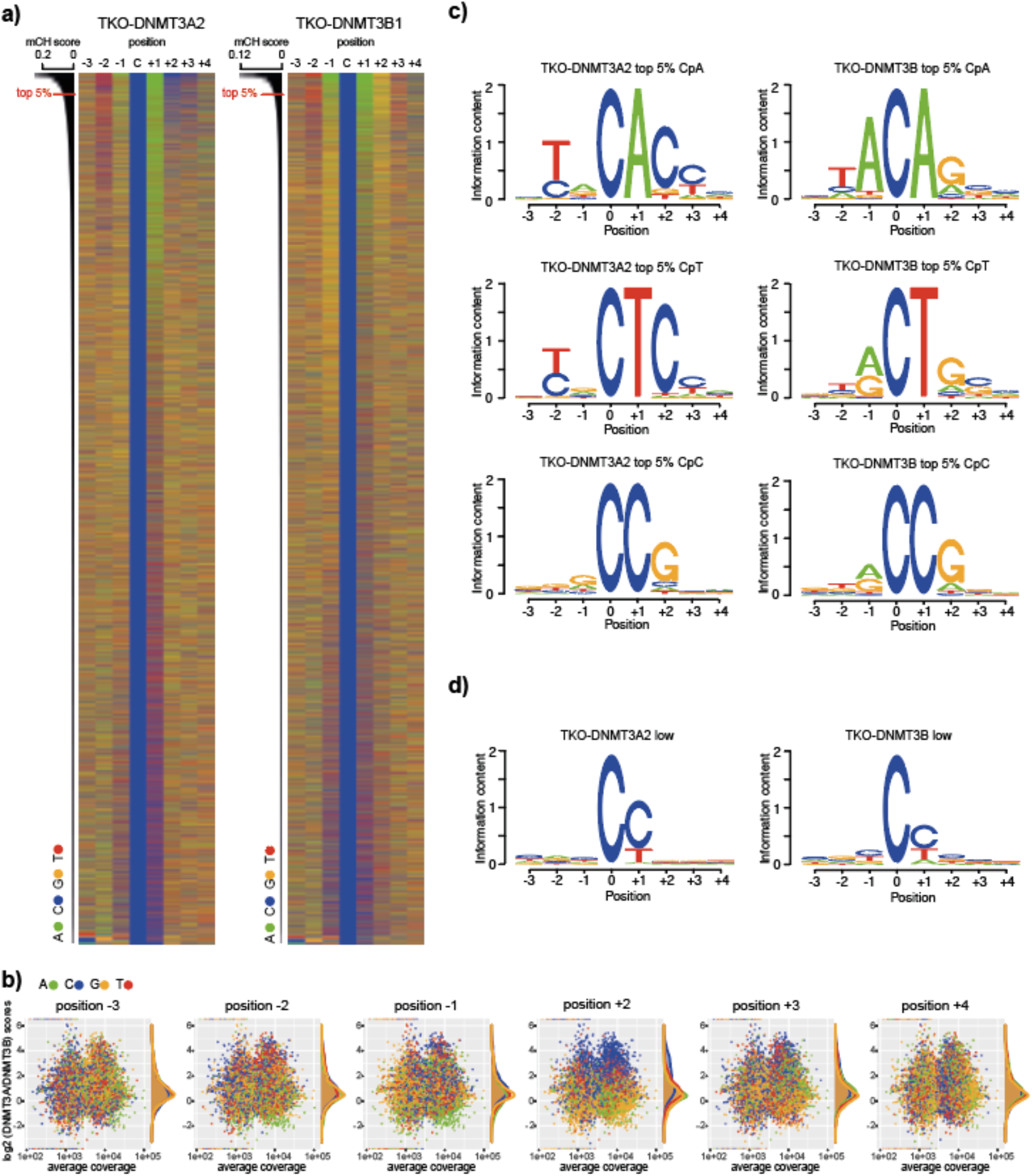
De novo DNMTs sequence preferences at non-CpG sites. **a)** Heatmap indicating the sequence composition at 8-mers ranked by CpH methylation for TKO-DNMT3A2 or TKO-DNMT3B1. Shown are the nucleotides at positions −2 to +4 around the methylated C. Green: A, blue: C, orange: G and red: T. The average methylation calculated for all instances of each 8-mer is shown above. **b)** MA plots showing log2 methylation differences between DNMT3A2 and DNMT3B1 activity at all analysed CpH 8-mers (y-axis) and the log10 average coverage of each 8-mer (x-axis). All plots show the same data, but the 8-mers are coloured based on the respective nucleotide at positions −3 to +4, excluding C and A positions 0 and +1. **c)** PWM obtained from top 5%-methylated CpH sites, calculated separately for CpA, CpT and CpC sites. **d)** PWM obtained from the least-methylated CpH sites.

**Supplementary Figure 6.**
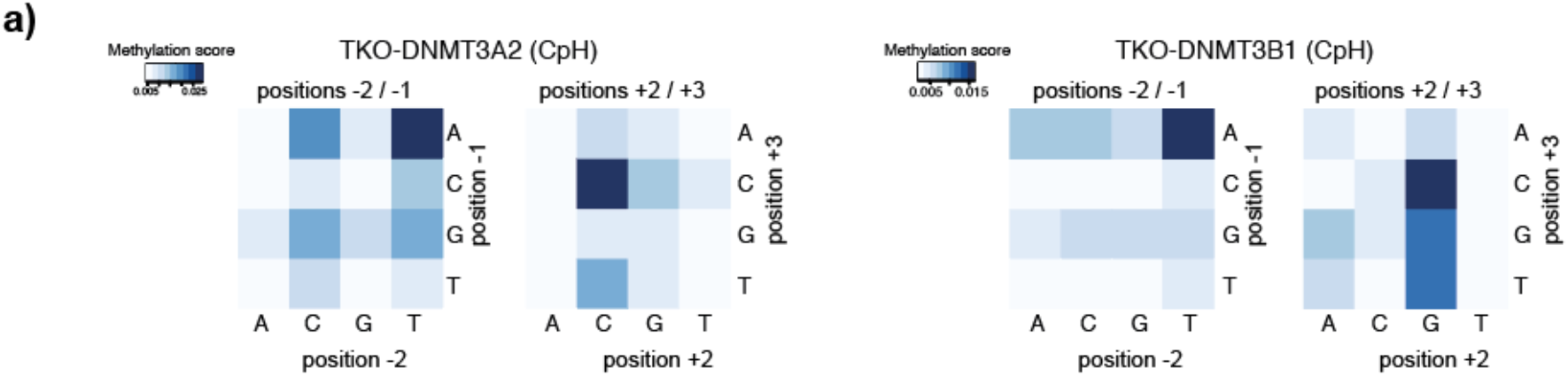
Nucleotide coupling at CpH flanking sequences. **a)** Heatmap showing the effect of dinucleotide coupling at positions −2/−1 or +2/+3 on CpH methylation (at position 0/+1). Shown is the methylation score at the cytosine in position 0, according to the indicated dinucleotide combinations, upstream or downstream.

**Supplementary Figure 7.**
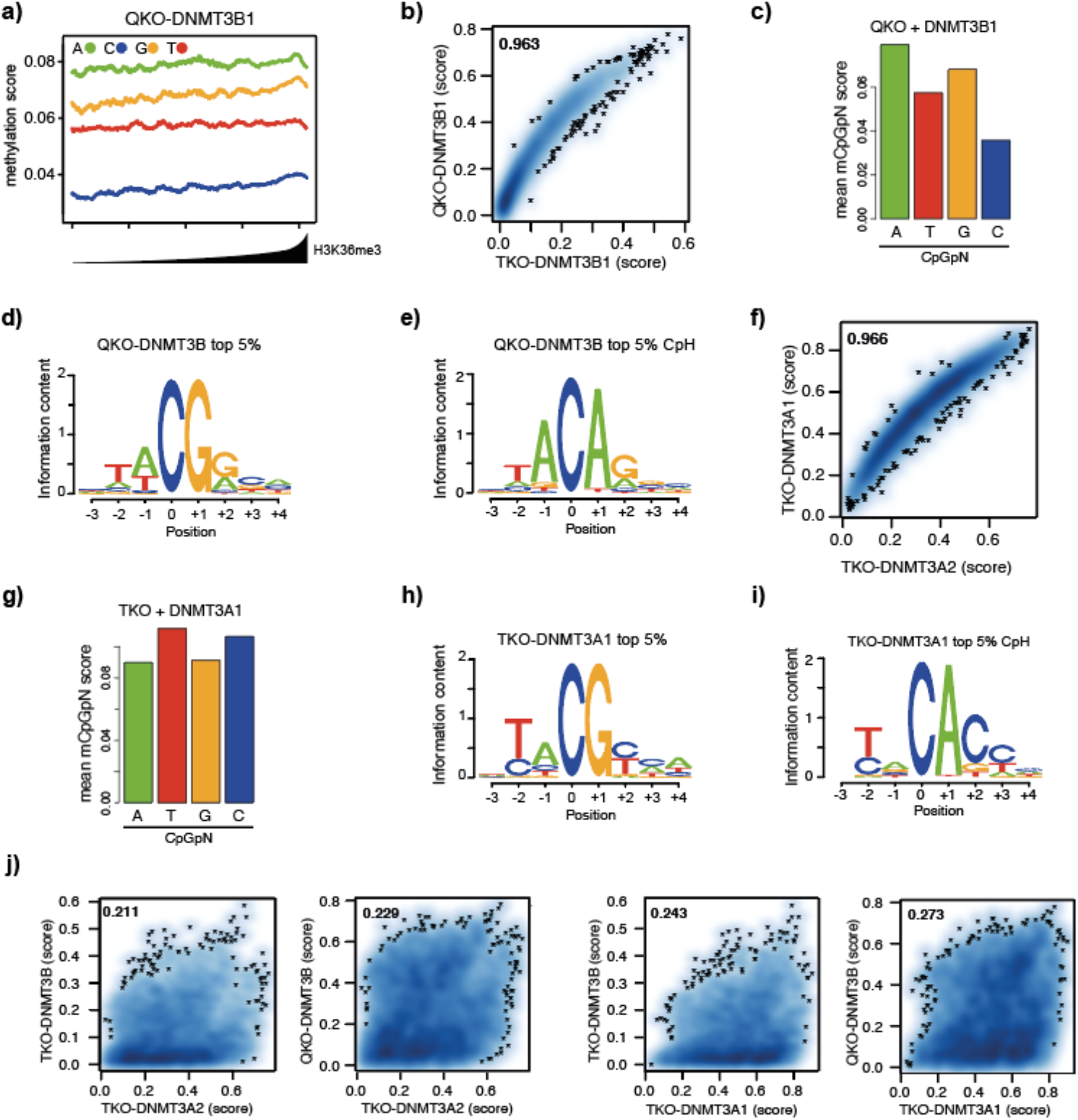
Differential genomic binding of DNMTs does not influence flanking sequence preference. **a)** Preferential de novo methylation of purines by DNMT3B is not altered in absence of H3K36me3. Shown are de novo DNA methylation at all four CpGpN context genome-wide in relation to H3K36me3 enrichment in wild type cells. 1-kb-sized genomic intervals were ranked and grouped by H3K36me3 enrichment (1000 intervals per bin) and DNA methylation was calculated per bin. Lines indicate mean DNA methylation per bin in TKO lacking H3K36me3 cells expressing DNMT3B1 for each CpGpN context. **b)** Scatter plot showing similarity in DNA methylation preferences between DNMT3B1 in absence or presence of H3K36me3 (QKO) at all 8-mers. **c)** Bar plots indicating average methylation scores at CpG sites followed by A, T, C and Gs (CpGpN) in the QKO-DNMT3B1 sample. **d-e)** Position weight matrix calculated from the top 5% 8-mers methylated by DNMT3B in absence of H3K36me3 indicate the preferred sequence at CpG (shown in d) and non-CpG sites (shown in e). **f)** Scatter plot showing similarity in DNA methylation preferences for all 8-mers between both DNMT3A isoforms. **g)** Bar plots indicating average methylation scores at CpG sites followed by A, T, C and Gs (CpGpN) in the TKO-DNMT3A1 sample. **h-i)** Position weight matrix calculated from the top 5% 8-mers methylated by DNMT3A1 indicate the preferred sequence at CpG (shown in h) and non-CpG sites (shown in i). **j)** Direct comparison between all DNMT3A and DNMT3B samples show low similarity in sequence preference (compared to b or f).

**Supplementary Figure 8.**
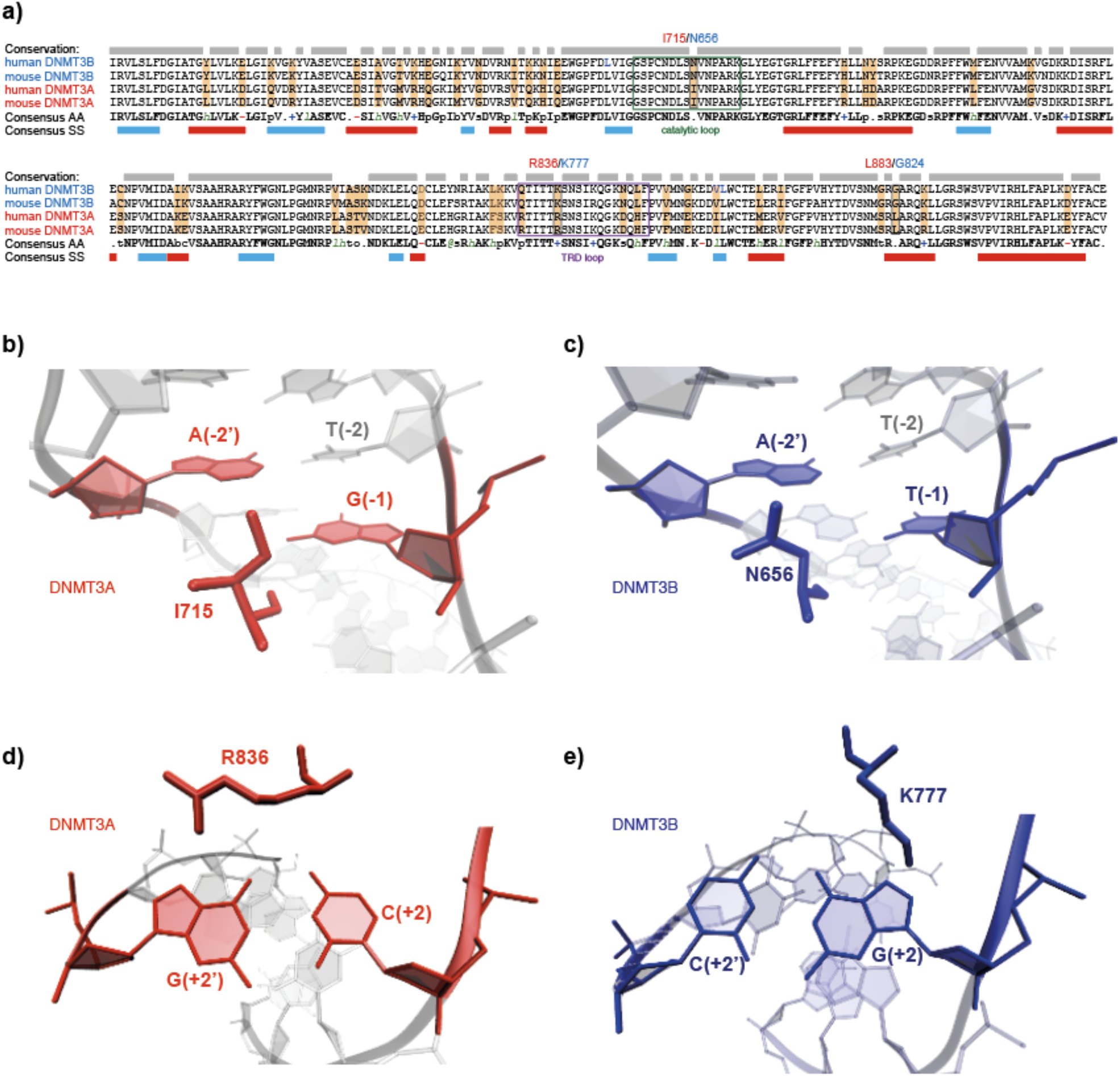
Structural differences between DNMT3A and DNMT3B provide potential cues for observed sequence preferences. **a)** Comparison of amino acid sequences of mouse/human DNMT3A and DNMT3B catalytic domains using PROMALS3D. Conservation is highlighted by grey boxes above, amino acid (AA) and secondary structure (SS) consensus is shown below. Conserved amino acids are indicated in bold, consensus secondary structures are indicated in red: a-helix or blue: b-sheet. Catalytic loop and TRD loops are highlighted by green or violet rectangles, respectively. Variant residues between DNMT3A and DNMT3B but conserved between the homologous proteins in human and mouse are highlighted in orange. Variant residues that interact with DNA based on DNAproDB using PDB ID: 6F57 are highlighted by black rectangles and their position in DNMT3A/DNMT3B is indicated above. **b-e)** Interactions between the variant residues DNMT3A-I715 / DNMT3B-N656 and DNMT3A-R836 / DNMT3B-K777 with DNA. Based on PDB IDs: 6F57 (DNMT3A) and 6KDA (DNMT3B).

**Supplementary Figure 9.**
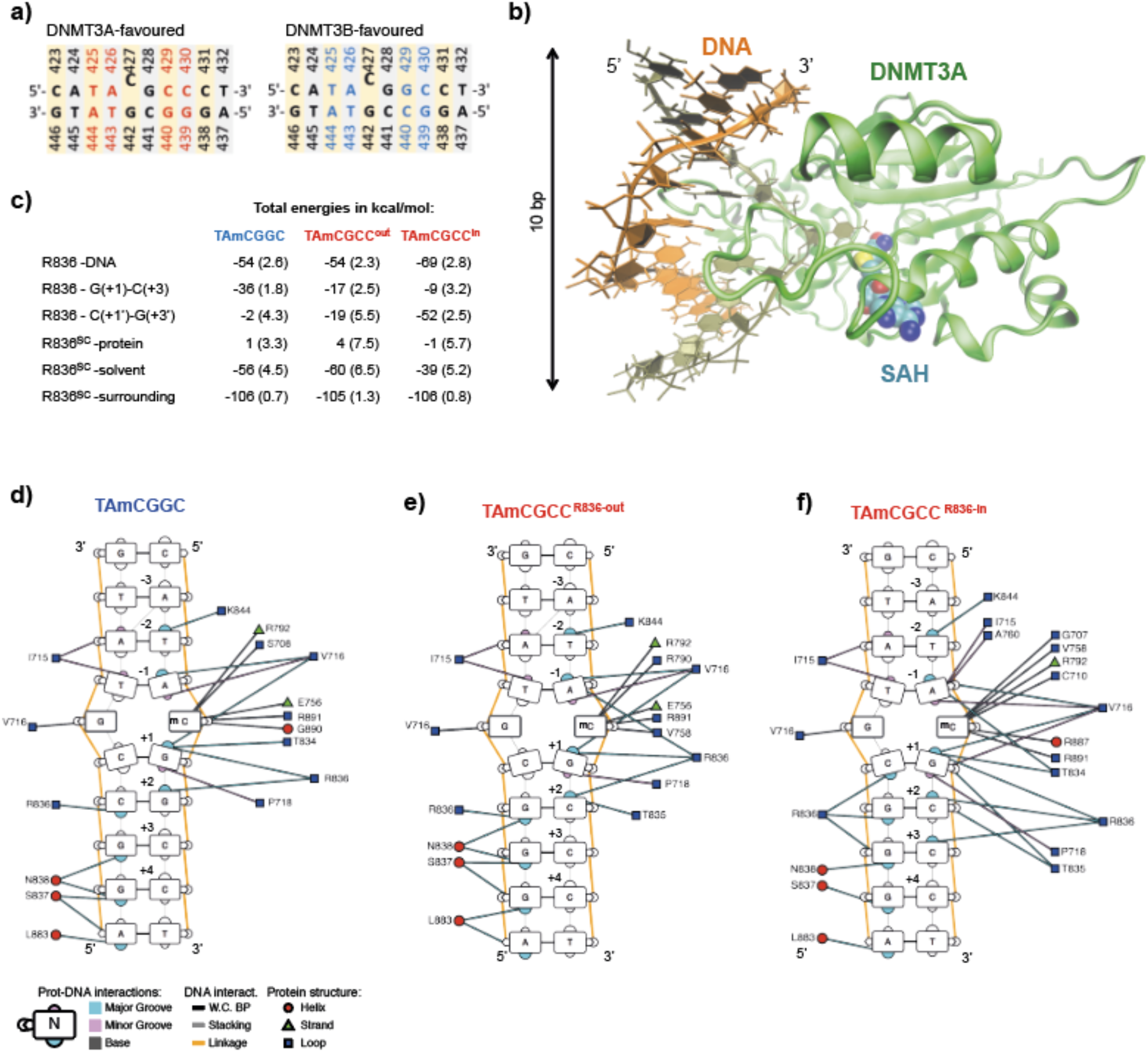
Molecular dynamics simulations reveal potential mechanisms underlying DNMT3A preference for CpGpC sites. **a)** The two DNA sequences used in the simulations with DNMT3A. The methylated cytosine (427 according to the numbering in the crystal structure 5YX2) is highlighted by the slight elevation. The nucleotides that were modified with respect to the DNA sequence in the crystal structure are shown in red (sequence preferred by DNMT3A) and blue (sequence preferred by DNMT3B). These two sequences differ only at position +2 (429) and the related base (440) in the base pair. **b)** Overview of the input structure (derived from PDB ID: 5YX2) that was used in the MD simulations. Green: DNMT3A, orange: methylated DNA (residues:C423-C432/G437-G446). The co-product of the reaction, S-adenosyl-L-homocysteine (SAH) is shown (spheres). **c)** Interaction energy between R836 and other components of the simulation systems. Each entry in the table is the sum of van der Waals and Coulombic terms. The statistical error (in parentheses) is calculated as the standard deviation of the mean over the five independent MD runs. **d-f)** Schematic overview of DNAproDB-reported intermolecular interactions between DNMT3A and DNA obtained from the MD simulations. Shown are interactions with disfavored (in d) and favored (in e and f) DNA duplex sequence. For the favored sequence, two schematics are shown corresponding to the state before the conformational switch (“R836-out” state, in e) and after (“R836-in” state, in f). Edges represent interactions between DNMT3A amino acid residues and DNA, while colours represent interactions with the major groove in blue, minor groove in pink and bases in grey. Position of interacting amino acids within DNMT3 protein secondary structures are shown as red circles for alpha helices, green triangles for beta strands and blue squares for loops.

